# Fpk1 regulates Cdr1 expression and ergosterol homeostasis in *Nakaseomyces glabratus* (*Candida glabrata*) during azole exposure

**DOI:** 10.64898/2026.08.14.744811

**Authors:** Samuel Cobb, Chelsea Chanheng, Cole Brown, Dawson Otey, Josie McFarland, Bao G. Vu

## Abstract

Azoles remain the most common antifungal therapy worldwide. However, *Nakaseomyces glabratus* (previously named *Candida glabrata*) has a high intrinsic tolerance against azole drugs. The organism can also accrue additional chromosomal mutations to elevate its resistant level during treatment. These genetic alterations often result in overexpression of the ABC transmembrane transporter Cdr1, which has been shown to directly transport drugs out of the fungal cells. Another resistant mechanism is the upregulation of the ergosterol biosynthesis pathway, which is the direct target of azoles. Although the mechanisms of azole resistance in *N. glabratus* are well defined, knowledge of their regulation remains limited. Here, we show that the protein kinase Fpk1 is required for optimal azole response *in vitro* and in an *in vivo* mouse infection model. Loss of Fpk1 gene or its kinase function significantly enhances azole sensitivity in both azole-susceptible and -resistant clinical isolates. Fpk1 function is required for optimal expression of Cdr1 upon azole challenge. It also influences the intracellular trafficking of ergosterol, without affecting its biosynthesis. Together, our data demonstrates the important role of Fpk1 function in the *N. glabratus* azole response and characterizes it as a new regulator of the efflux pump and ergosterol biosynthesis pathways.

**IMPORTANCE:** Antifungal treatment against life-threatening bloodstream *Candida* infection remains limited to azoles, echinocandins, and polyenes. Among them, azoles are the most prescribed therapy worldwide. However, the pathogenic yeast *Nakaseomyces glabrataus* has a high level of resistance against azoles (> 10%) (1). This often complicates treatment and increases mortality and morbidity rates. Therefore, understanding the mechanism of azole resistance would reinforce the treatment strategy and bolster future therapy development. Here, we identify the protein kinase Fpk1 as an important regulator of the drug efflux plump and ergosterol biosynthesis pathways. Disruption of the Fpk1 function significantly enhances the azole efficacy *in vitro* and in a mouse model of *Candida* systemic infection. Protein kinases are druggable targets, and our data presents Fpk1 as a viable candidate for future antifungal development.

## OBSERVATION

*Nakaseomyces glabratus* (aka. *Candida glabrata*) is the second leading cause of *Candida* bloodstream infection in the US (1, 2). The organism has high intrinsic tolerance against azole drugs (the most common antifungal treatment worldwide) and can readily acquire additional mutations to become hyper-resistant during treatment (3–5). Recently, our lab has constructed a protein kinase (PK) deletion library in *N. glabratus* and examined the PK functions in drug resistance. A thorough report of this data will be presented elsewhere, but one of the PKs of interest that exhibited a significant role in azole resistance was the flipase kinase 1 (Fpk1). In *Saccharomyces cerevisiae*, Fpk1 and its homolog Fpk2 regulate cell membrane phospholipid asymmetry through maintaining the function of Dnf1 and Dnf2 flippases (6, 7). Fpk1/2 kinase activity is also important in maintaining intracellular sphingoid long-chain base levels, which are building blocks of membrane sphingolipids (8). Together with ergosterol, phospholipids, and sphingolipids are essential components of the fungal cell membrane (9, 10).

Azoles target fungal cell membrane homeostasis by inhibiting the biosynthesis of ergosterol (4). However, the impact of Fpk1/2 on modulating the azole effect in fungi remains unexplored. In *N. glabratus*, there is only copy of the Fpk gene, *FPK1* (CAGL0C03509g) and, in this study, we provide evidence that Fpk1 is required for *N. glabratus* optimal azole resistance.

To examine the role of Fpk1 in antifungal resistance, we first constructed an *FPK1* isogenic deletion (*fpk1*Δ) in CBS138 wildtype (WT) and azole-resistant backgrounds carrying a *PDR1* gene mutant allele, D1082G or R376W (11). Spot-test assays were performed with fluconazole, micafungin, and amphotericin B. The loss of the *FPK1* significantly increased fluconazole sensitivity in both the WT and azole resistant isolates (Figure 1A,1B). This effect seemed to be specific to fluconazole as there was no change in the micafungin response and a slight increase in amphotericin B resistance among the *fpk1Δ* mutants (Figure 1A,1B). Chromosomal complementation of *FPK1* at the native locus recovered azole tolerance to the WT level, while the kinase-dead (KD) allele (*FPK1_D622A_*) complementation behaved like the null-mutant, suggesting that the kinase activity of Fpk1 was responsible for the observed phenotype (Supplemental figure 1A). To rule out a strain-dependent phenotype, we also confirmed the enhanced azole sensitivity of *fpk1Δ* mutants in SM1 and SM3 backgrounds (Supplemental figure 1B).

**Figure 1.**
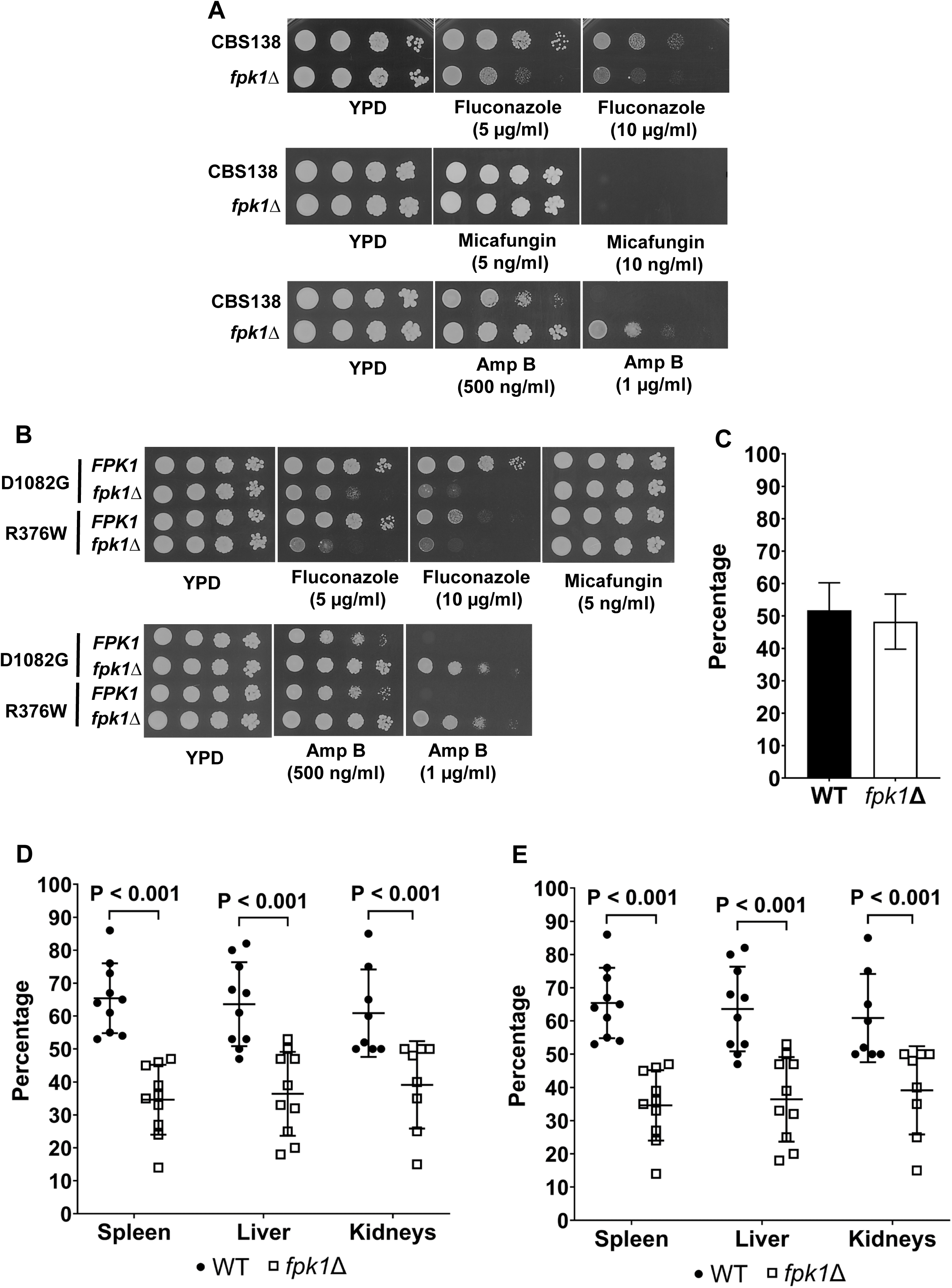
Mid-log cells were 10-fold serially diluted and spotted onto solid YPD agar containing various concentrations of fluconazole, micafungin, and amphotericin B (Amp B). (A) Spot-test assay of CBS138 wildtype and its *FPK1* gene isogenic deletion (*fpk1*Δ) counterpart, and (B) CBS138 isolates carrying *PDR1* gene allele D1082G or R36W and their *FPK1* gene isogenic deletion (*fpk1*Δ) counterparts. In separate experiments, mice were challenged with 1:1 mixture of WT-mCherry and *fpk1Δ* isolates, treated with posaconazole or saline for 8 days, and fungal colony forming units (CFUs) were assessed in various organs. (C) The starting inoculum of WT-mCherry and *fpk1Δ* isolates. (D) CFU distribution between the two isolates in spleen, liver, and kidneys without treatment, and (E) with posaconazole treatment. Statistical differences between indicated groups were performed by unpaired T-test analysis.

Next, we expanded our analysis to other azoles and echinocandins by performing a broth microdilution assay with fluconazole, voriconazole, itraconazole, posaconazole, caspofungin and micafungin. Being consistent with the spot-test assay, a 2-fold reduction in MIC_50_ of all tested azoles was observed among *fpk1Δ* mutants in the CBS138 WT background, while there was no change in echinocandin MIC_50_ (Supplemental table 1). Together, these data have indicated that Fpk1 is important in maintaining azole tolerance (in CBS138 and SM1 isolates) and resistance (in *PDR1* mutant and SM3 isolates) levels in *N. glabratus*.

Having established the role of Fpk1 *in vitro*, we analyzed its effect *in vivo* using a mouse systemic infection model. We first generated a WT-mCherry strain from the CBS138 and challenged (via tail-vein injection) mice with a 1:1 mixture of WT-mCherry and *fpk1Δ* isolates. Mice were then treated with posaconazole or saline for 8 days, and fungal colony forming units (CFUs) were assessed in kidneys, spleens, and livers. Even without azole treatment, the WT-mCherry outgrew the *fpk1Δ* in all three organs, suggesting a role of Fpk1 in fungal *in vivo* fitness (Figure 1C, 1D). With posaconazole treatment, *fpk1Δ* was further outcompeted by the WT-mCherry, especially in the kidney with a 9:1 (WT-mCherry: *fpk1Δ*) distribution (Figure 1E).

To rule out any confounding effect from the mCherry expression cassette, in separate experiments, we challenged mice with a 1:1 mixture of WT and *fpk1Δ*-mCherry isolates. Consistently, *fpk1Δ*-mCherry was significantly outcompeted by the WT in the mouse kidneys after 8 days of posaconazole treatment (supplemental figure 2A, 2B). These data have shown that Fpk1 is also required for *N. glabratus* optimal azole tolerance *in vivo*.

In *N. glabratus*, azole tolerance can arise from an overexpression of the ABC transporter Cdr1 or upregulation of the ergosterol biosynthesis (ERG) pathway (12). Next, we examined the effect of Fpk1 on Cdr1 level. WT, *fpk1Δ*, *FPK1* complement (*fpk1Δ*::FPK1) and *FPK1*-KD isolates were treated with fluconazole or a vehicle for 3 hours. Total protein was extracted and Cdr1 was examined by Western immunoblotting with a rabbit anti-Cdr1 antibody. Specificity of the primary antibody was verified (supplemental figure 3). Although there was no difference in the basal expression of Cdr1 among tested isolates, *fpk1Δ* and *FPK1*-KD failed to induce Cdr1 expression to the same level as WT and *FPK1* complement (*fpk1Δ*::*FPK1*) isolates with azole treatment (Figure 2A, 2B). These data have suggested that the Fpk1 kinase activity is required for the induction of Cdr1 by fluconazole which, in turn, supports the increased azole sensitivity among *fpk1Δ* mutants.

**Figure 2.**
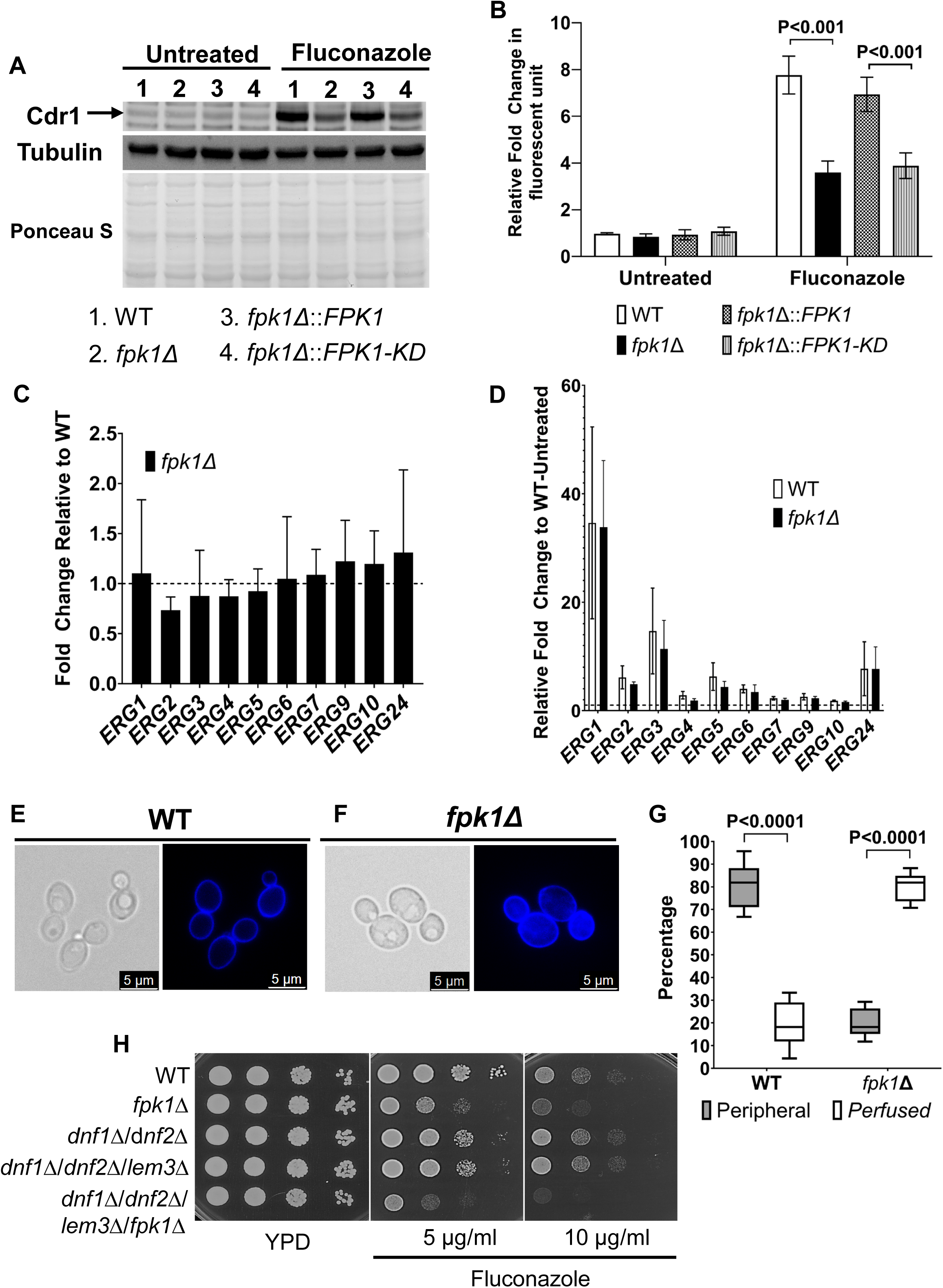
The effects of Fpk1 on Cdr1 and ERG gene expression were assessed. (A) Western immunoblot analysis of Cdr1 protein expression with and without fluconazole treatment in WT, *fpk1Δ*, *FPK1* WT complement (*fpk1Δ*::*FPK1*), and *FPK1*-Kinase Dead complement (*FPK1*-KD) isolates using a rabbit polyclonal antibody raised against Cdr1. Ponceau S and Tubulin were used as loading controls. (B) Quantification of the band intensity from Figure 1A from three biological replicates. Statistical differences between indicated groups were performed by unpaired T-test analysis. (C) RT-qPCR analyses of representative ERG gene expression in WT and *fpk1Δ* without fluconazole treatment, and (D) with treatment; dotted line represents baseline of untreated WT. In separate experiments, mid-log, live WT and *fpk1Δ* cells were labeled with filipin dye and observed under a confocal microscope. Representative of phase contrast and filipin staining images of (E) WT and (D) *fpk1Δ* cells. Uncropped images are presented in Supplemental Figure 4 and 5. (G) Quantification of “peripheral” and “perfused” phenotype in WT and *fpk1Δ* cells. Statistical differences between indicated groups were performed by paired T-test analysis. (H) mid-log WT, *fpk1Δ*, *dnf1*Δ/*dnf2*Δ (double deletion), *dnf1*Δ/*dnf2*Δ/*lem3*Δ (triple deletion), and *dnf1*Δ/*dnf2*Δ/*lem3*Δ/*fpk1Δ* (quadruple deletion) cells were 10-fold serially diluted and spotted onto solid YPD agar containing various concentrations of fluconazole.

To assess Fpk1 function in the ergosterol biosynthesis pathway, we compared the expression levels of representative ERG genes in the WT and *fpk1Δ* by RT-qPCR. However, there was no significant difference in ERG gene expression between the two groups regardless of azole treatment (Figure 2C, 2D). Next, we investigated the distribution of ergosterol in the cell. Mid-log, live WT and *fpk1Δ* cells were labeled with filipin (an ergosterol stain) and immediately observed under a confocal microscope. While filipin was distributed evenly on the WT cell surface (we labeled this phenotype as “peripheral”), among *fpk1Δ* isolates filipin was perfused throughout the cytoplasm (we labeled this phenotype as “perfused”) (Figure 2E, 2F, 2G). This has suggested a dysregulation of ergosterol trafficking in *fpk1Δ* background. Together, these outcomes have indicated that Fpk1 has an impact on intracellular ergosterol distribution instead of its biosynthesis.

Since Fpk1 has been shown to regulate the activity of Dnf1 and Dnf2 flippases (6), we tested the impact of these gene deletion on azole sensitivity. We also included Lem3 in the analysis, since it is an essential adaptor protein of Dnf1 and Dnf2 (6). Interestingly, double-deletion of Dnf1 and Dnf2 (*dnf1*Δ/*dnf2*Δ) and triple deletion of all three genes (*dnf1*Δ/*dnf2*Δ/*lem3*Δ) shows no significant change in fluconazole response relative to the WT in a spot-test assay (Figure 2H). However, deletion of *FPK1* together with the flippase complex (*dnf1*Δ/*dnf2*Δ/*lem3*Δ/*fpk1Δ*) sensitized the cell toward fluconazole even more than *fpk1*Δ alone (Figure 2H). These data have suggested that the observed effect of Fpk1 on azole sensitivity is independent of the flippase activity. However, further assessing this novel mechanism of Fpk1 is beyond the scope of this study.

Together, these findings are consistent with Fpk1 acting as an upstream modulator of azole susceptibility in both azole-sensitive and -resistant isolates. The known mechanism of Fpk1 activating Dnf1 and Dnf2, flippase seems to be independent to this novel effect on azole sensitivity since *dnf1*Δ/*dnf2*Δ/*lem3*Δ exhibits no change in azole response. Fpk1 kinase activity is required for the induction of the efflux-pump Cdr1 with azole treatment. Additionally, the loss of Fpk1 function promotes intracellular accumulation of ergosterol without any apparent effect on its biosynthesis. While both mechanisms provide explanation for enhanced azole susceptibility, intracellular accumulation of ergosterol also illuminates the slight increase in amphotericin B tolerance among *fpk1Δ* mutants. Collectively, this study identifies Fpk1 as a regulator of Cdr1 and provides a new mechanism of inhibiting ergosterol function through disrupting its intracellular distribution.

## MATERIALS AND METHODS

### Strains and growth conditions

*N. glabratus* was grown in rich YPD medium [1% yeast extract, 2% peptone, 2% glucose]. All solid media contained 1.5% agar. Nourseothricin (Gold Biotechnology, Olivette, MO) was supplemented to YPD media at 60 μg/ml to select strains containing nourseothricin expressing cassette (*NAT*). All strains used in this study are listed in Supplemental Table 1.

### Plasmid and yeast transformation

Non-autonomous plasmids were constructed on the pUC19 background (New England Biolabs, Ipswitch, MA). All isogenic deletion constructs were made by assembling the recyclable cassette from pBV65 (13) and fragments from the immediate upstream/ downstream regions of the target genes. Gibson assembly cloning (New England Biolabs, Ipswich, MA) was employed to assemble fragments together. Eviction of the recyclable cassette left a single copy of *LOXP* in place of the excised target gene coding region. The recycling *NAT* cassette was detailed elsewhere (13). Gene complementation was built from isogenic deletion backgrounds, and the same strategy (as gene deletion) was used with a new inserting cassette.

The kinase dead mutant *FPK1_D622A_* was generated by PCR amplification of two fragments of the *FPK1* coding sequence along the kinase domain using primers that contained a point mutation on the overhang. The point mutation was generated in the homologous locus to the active kinase site described in *Saccharomyces cerevisiae FPK1* (6). These fragments were annealed via Gibson assembly (New England Biolabs) alongside the plasmid backbone and a selection cassette.

Transformations were performed using a lithium acetate method (14). After heat shocking, cells were allowed to incubate statically overnight at 30°C, before being plated onto YPD agar plates supplemented with 60 µg/mL nourseothricin and/or 2mM methionine. Plates were incubated at 30°C for 48 hours. Individual colonies were isolated and screened by PCR to ensure correct insertion of the construct into the chromosome.

### Broth microdilution assay

Broth microdilution assays were performed based on the Clinical and Laboratory Standards Institute (CLSI) guidelines. In short, clear, round bottom 96-well plates (MIDSCI, Fenton, MO) were used. 10^3^ mid-log cells were used as inoculum. Fluconazole (1-256 µg/mL), voriconazole (0.03125-8 µg/mL), itraconazole (0.03125-8 µg/mL), posaconazole (0.015265-4 µg/mL), caspofungin (4-1024 ng/mL), micafungin (1- 256ng/mL), and amphotericin B (8-2048ng/mL) were tested. Plates were incubated at 37° for 24 hours before being analyzed visually. The MIC_50_ determined at which drug concentration growth was reduced by approximately 50%.

### Spot test assay

Mid-log cultures were diluted in PBS to an O.D. of 0.3. Cells were 10-fold serially diluted and spotted onto YPD agar plates containing different concentrations of fluconazole, micafungin, and amphotericin B. Plates were incubated at 37°C for 24 hours before imaging was performed.

### Quantification of transcript levels by RT-qPCR

Mid-log cells were treated with the vehicle or fluconazole (60 µg/ml) for 3 hours in YPD at 37°C. Total RNA was extracted from cells by using TRIzol (Invitrogen, Carlsbad, CA) and chloroform (Fisher Scientific, Hampton, NH) followed by purification with an RNA mini kit (Invitrogen, Carlsbad, CA). Total RNA was reverse transcribed using an iScript cDNA synthesis kit (Bio-Rad, Des Plaines, IL). qPCR was performed with iTaq universal SYBR green supermix (Bio-Rad). Target gene transcript levels were normalized to transcript levels of 18S rRNA. Data reported are from three biological replicates.

### Filipin staining

Mid-log cells were washed twice with phosphate buffer saline (PBS). Cells were resuspended in 1ml of PBS and stained with 2 µg/ml of filipin (Tocris BioScience, Minneapolis, MN) for 30 minutes at 30°C, in the dark. Cells were then centrifuged at 3500 x g for 5 minutes, resuspended in 300 µl of PBS, and immediately observed under the Leica THUNDER DMi8 imager system (Leica, Boston, MA). Images were analyzed with the Leica LIGHTNING software (Leica, Boston, MA). Filipin was stocked at 2 mg/ml in DMSO. “Peripheral” and “Perfused” filipin staining phenotypes were analyzed and counted visually. A total of 500 cells were assessed in each isolate.

### Western immunoblotting

Harvested cells were washed twice with PBS and lysed with lysis buffer [1.85 M NaOH, 7.5% 2-Mercaptoethanol] on ice. Proteins were precipitated with 50% Trichloroacetic acid on ice and resuspended in sample buffer [40 mM Tris pH8, 8.0M Urea, 5% SDS, 1% 2-Mercaptoethanol]. Secondary antibodies (StarBright) were purchased from Bio-Rad (Des Plaines, IL. Imaging was performed using the BioRad ChemiDoc MP Imaging System (Bio-Rad) and analyzed by Image Studio Lite Software (LI-COR Biosciences, Lincoln, NE). Detected target band fluorescence intensity was normalized against tubulin fluorescence intensity and compiled from 3 biological replicates.

To generate the rabbit polyclonal antibody against Cdr1, the first 172 amino acids of Cdr1 protein sequence were cloned into pET28a+ (EMD Millipore Inc.) plasmid to generate *CDR1_172_-6xHIS*. Protein expression was carried out in *Escherichia coli* BL21(DE3) (Thermo Scientific, Waltham, MA). Transformants were grown to log phase and induced with 1 mM isopropyl-β-d-thiogalactopyranoside (IPTG) for 90 min at 37°C. Protein purification was accomplished using Talon metal affinity resin (TaKaRa Bio USA, Inc.) as described by the manufacturer. Purified proteins were then dialyzed against PBS, lyophilized, and sent to Pacific Immunology (Ramona, CA) for polyclonal antibody generation. Western immunoblot was performed with CBS138 (WT), CBS138/*cdr1*Δ, and fluconazole treated WT to test the antibody specificity (Supplemental figure 3).

### Animal studies

Around 1 × 10^8^ colony forming unit (CFU) of each tested isolates were combined to generate a 1:1 mixture inoculum. 100 µl of mixed culture was inoculated into CD-1 mice (Jackson Laboratory, Bar Harbor, ME) via tail-vein injection. 5 male and 5 female mice were tested in each group. In antifungal treated experiments, after 24-hour post-infection, mice were treated with posaconazole (5 mg/kg) or saline, via intraperitoneal injection, daily for 8 days. At the end of the experiment, mice were euthanized via CO_2_ asphyxiation. Kidneys, spleen, and liver were harvested and washed with sterile PBS. All organs were lysed using 70 µm cell strainers (VWR, Radnor, PA) in PBS. Both kidneys were combined during lysis. Lysate was serially diluted and plated onto YPD agar. Plates were incubated at 37°C for at least 24 hours before being evaluated. Data was reported as the percentage of normal and red colonies on the plate post-incubation. Individual mouse kidney data was reported as the average of the two kidneys harvested from each mouse.

All animal procedures were approved by the Institutional Animal Care and Use Committee at the University of Oklahoma Health Campus (OUHC) (protocol 23-075- CHI). All work was performed in accordance with the recommendations of the Guide for the Care and Use of Laboratory Animals of the National Institutes of Health.

### Statistics

Significance of results was determined using the T-test. Unpaired conditions were used to assess isolates with different genetic backgrounds or among outbred mice. Paired conditions were used to compare different treatment conditions (or analyses) on the same genetic background.

---

**Supplemental Figure 1.** Mid-log cells were 10-fold serially diluted and spotted onto solid YPD agar containing various concentrations of fluconazole. (A) Spot-test assay of CBS138 wildtype, its *FPK1* gene isogenic deletion (*fpk1*Δ), *FPK1* WT complement (*fpk1Δ*::*FPK1*), and *FPK1*-Kinase Dead (*FPK1*-KD) complement isolates. (B) Spot-test assay of wildtype SM1 and SM3 strains and their *FPK1* gene isogenic deletion (*fpk1*Δ) counterparts.

**Supplemental Figure 2.** Mice were challenged with 1:1 mixture of CBS138 (WT) and *fpk1Δ*-mCherry isolates, treated with posaconazole or saline for 8 days, and fungal colony forming units (CFUs) were assessed in kidneys. A) The starting inoculum of WT and *fpk1Δ*-mCherry isolates. (B) CFU distribution between the two isolates in kidneys with azole treatment. Statistical differences between indicated groups were performed by unpaired T-test analysis.

**Supplemental Figure 3.** Verification of rabbit polyclonal antibody raised against Cdr1 were tested by Western immunoblotting Cdr1 protein in CBS138 (WT), CBS138 *cdr1*Δ, and CBS138 treated with fluconazole. Ponceau S and Tubulin were used as loading controls.

**Supplemental Figure 4.** A representative image of filipin staining in CBS138 WT.

**Supplemental Figure 5.** A representative image of filipin staining in *fpk1*Δ mutant.

**Supplemental Table 1.** Strains used in this study.

| Strain | Genotype | Parent strain | Reference |
| --- | --- | --- | --- |
| CBS138 | Wildtype | N/A | ATCC2001 |
| CBS138, *fpk1*Δ | *fpk1*Δ::*LOXP* | CBS138 | This study |
| *FPK1* complement | *fpk1*Δ::*FPK1*::*LOXP* | *fpk1*Δ::*LOXP* | This study |
| *FPK1* kinase dead | *fpk1*Δ::*FPK1*<sub>D622A</sub>::*LOXP* | *fpk1*Δ::*LOXP* | This study |
| Pdr1-D1082G | *pdr1*Δ::*PDR1*<sub>D1082G</sub>::*LOXP*<br>*pdr1*Δ::*PDR1*<sub>D1082G</sub>::*LOXP* | CBS138, *pdr1*Δ | (11) |
| Pdr1-D1082G, *fpk1*Δ | *fpk1*Δ::*LOXP* | Pdr1-D1082G | This study |
| Pdr1-R376W | *pdr1*Δ::*PDR1*<sub>R376W</sub>::*LOXP*<br>*pdr1*Δ::*PDR1*<sub>R376W</sub>::*LOXP* | CBS138, *pdr1*Δ | (11) |
| Pdr1-R376W, *fpk1*Δ | *fpk1*Δ::*LOXP* | Pdr1-R376W | This study |
| SM1 | Wildtype | N/A | (15) |
| SM3 | Wildtype | N/A | (15) |
| SM1, *fpk1*Δ | *fpk1*Δ::*LOXP* | SM1 | This study |
| SM3, *fpk1*Δ | *fpk1*Δ::LOXP | SM3 | This study |
|  | *dnf1*Δ::LOXP |  |  |
| *dnf1*Δ/*dnf2*Δ | *dnf2*Δ::LOXP | CBS138 | This study |
|  | *dnf1*Δ::LOXP |  |  |
|  | *dnf2*Δ::LOXP |  |  |
| *dnf1*Δ/*dnf2*Δ/*lem3*Δ | *lem3*Δ::LOXP | CBS138 | This study |
|  | *dnf1*Δ::LOXP |  |  |
|  | *dnf2*Δ::LOXP |  |  |
|  | *lem3*Δ::LOXP |  |  |
| *dnf1*Δ/*dnf2*Δ/*lem3*Δ/*fpk1*Δ | *fpk1*Δ::LOXP | CBS138 | This study |

**Supplemental Table 2.** Minimum inhibitory concentration (MIC) of fluconazole, voriconazole, itraconazole, posaconazole, caspofungin and micafungin in CBS138 (WT) and its *fpk1Δ* counterpart.

|  | Fluconazole<br>(μg/mL) | Voriconazole<br>(μg/mL) | Itraconazole<br>(μg/mL) | Posaconazole<br>(μg/mL) | Caspofungin<br>(ng/mL) | Micafungin<br>(ng/mL) |
| --- | --- | --- | --- | --- | --- | --- |
| **WT** | 16 | 0.5 | 0.5 | 1 | 32 | 16 |
| ***fpk1*Δ** | 8 | 0.25 | 0.25 | 0.5 | 32 | 16 |

## REFERENCES.

1. Beardsley J, Kim HY, Dao A, Kidd S, Alastruey-Izquierdo A, Sorrell TC, Tacconelli E, Chakrabarti A, Harrison TS, Bongomin F, Gigante V, Galas M, Siswanto S, Dagne DA, Roitberg F, Sati H, Morrissey CO, Alffenaar JW. 2024. Candida glabrata (Nakaseomyces glabrata): A systematic review of clinical and microbiological data from 2011 to 2021 to inform the World Health Organization Fungal Priority Pathogens List. Med Mycol 62.

2. Bays DJ, Jenkins EN, Lyman M, Chiller T, Strong N, Ostrosky-Zeichner L, Hoenigl M, Pappas PG, Thompson Iii GR. 2024. Epidemiology of Invasive Candidiasis. Clin Epidemiol 16:549–566.

3. Naskar S, Prajapati A, Kaur R. 2025. Antifungal drug resistance in Candida glabrata: role of cellular signaling and gene regulatory networks. FEMS Yeast Res 25.

4. Li Y, Hind C, Furner-Pardoe J, Sutton JM, Rahman KM. 2025. Understanding the mechanisms of resistance to azole antifungals in Candida species. JAC Antimicrob Resist 7:dlaf106.

5. de Oliveira Santos GC, Vasconcelos CC, Lopes AJO, de Sousa Cartagenes MDS, Filho A, do Nascimento FRF, Ramos RM, Pires E, de Andrade MS, Rocha FMG, de Andrade Monteiro C. 2018. Candida Infections and Therapeutic Strategies: Mechanisms of Action for Traditional and Alternative Agents. Front Microbiol 9:1351.

6. Roelants FM, Baltz AG, Trott AE, Fereres S, Thorner J. 2010. A protein kinase network regulates the function of aminophospholipid flippases. Proc Natl Acad Sci U S A 107:34–9.

7. Nakano K, Yamamoto T, Kishimoto T, Noji T, Tanaka K. 2008. Protein kinases Fpk1p and Fpk2p are novel regulators of phospholipid asymmetry. Mol Biol Cell 19:1783–97.

8. Yamane-Sando Y, Shimobayashi E, Shimobayashi M, Kozutsumi Y, Oka S, Takematsu H. 2014. Fpk1/2 kinases regulate cellular sphingoid long-chain base abundance and alter cellular resistance to LCB elevation or depletion. Microbiologyopen 3:196–212.

9. Parks LW, Casey WM. 1995. Physiological implications of sterol biosynthesis in yeast. Annu Rev Microbiol 49:95–116.

10. Mbuyane LL, Bauer FF, Divol B. 2021. The metabolism of lipids in yeasts and applications in oenology. Food Res Int 141:110142.

11. Simonicova L, Moye-Rowley WS. 2020. Functional information from clinically-derived drug resistant forms of the Candida glabrata Pdr1 transcription factor. PLoS Genet 16:e1009005.

12. Vu BG, Thomas GH, Moye-Rowley WS. 2019. Evidence that Ergosterol Biosynthesis Modulates Activity of the Pdr1 Transcription Factor in Candida glabrata. mBio 10.

13. Vu BG, Moye-Rowley WS. 2018. Construction and Use of a Recyclable Marker To Examine the Role of Major Facilitator Superfamily Protein Members in Candida glabrata Drug Resistance Phenotypes. mSphere 3.

14. Kawai S, Hashimoto W, Murata K. 2010. Transformation of Saccharomyces cerevisiae and other fungi: methods and possible underlying mechanism. Bioeng Bugs 1:395–403.

15. Magill SS, Shields C, Sears CL, Choti M, Merz WG. 2006. Triazole cross-resistance among Candida spp.: case report, occurrence among bloodstream isolates, and implications for antifungal therapy. J Clin Microbiol 44:529–35.

